# Who, what, when, where: Phenological trends of the Seedcorn Maggot Complex (Diptera: Anthomyiidae) in three vegetable crops in Southern Québec

**DOI:** 10.64898/2026.05.22.725509

**Authors:** Allen Bush-Beaupré, Anne-Marie Fortier, Jade Savage

## Abstract

In eastern Canada, the seedcorn maggot complex (SMC) includes three agriculturally important taxa: *Delia platura* biotypes H and N, and *D. florilega*. Because their larvae are morphologically indistinguishable, field-collected individuals are often grouped together despite differences in phenology and life history. Using larval collections from broccoli, green onion, and dry onion fields in Southern Québec (2017–2022), we identified specimens via HRM analysis and characterized taxon-specific phenology and abundance using Bayesian hierarchical generalized additive models. In *Allium* crops, *D. platura* biotype N dominated early-season infestations (May), followed by a shift in early June towards increased abundance of biotype H and *D. florilega*. In broccoli, all SMC members appeared later, with overlapping abundance peaks in June. These results reveal strong crop-dependent phenological differences and underscore the need to treat SMC members as distinct biological entities in pest management. The implications for optimizing an emerging sterile insect technique (SIT) program for *D. platura* in Québec are significant: field releases of sterile males should target early-season biotype N in *Allium* crops, followed by releases of both biotypes later in the season in broccoli to maximize control efficacy.

## Introduction

In eastern Canada, the seedcorn maggot complex (SMC) counts three biological entities of agriculture relevance, namely the seedcorn maggot, *Delia platura* (Meigen) biotypes H and N, and the bean seed maggot, *D. florilega* (Zetterstedt) (Bush-Beaupré et al., 2024; Griffiths, 1993; Savage et al., 2016). Most life stages are morphologically identical for these three taxa except for male *D. florilega* who display diagnostic leg chaetotaxy (Savage et al. 2016). Being highly polyphagous, SMC larvae feed on many host plant species including cultivated vegetables and field crops (Griffiths, 1993; Hough-Goldstein & Hess, 1984; Howard et al., 1994; Soroka & Dosdall, 2011) and depending on locality, *D. florilega* is generally reported in lower relative abundance than either (or both) biotypes of *D. platura* in the majority of crops it infests in Canada (Griffiths, 1993; Miller & McClanahan, 1960; Savage et al., 2016).

Reports on the relative contribution to damage caused to vegetable crops by each SMC member in the field and details pertaining to host associations remain unclear in North America as molecular identification of identical SMC larvae (including those of the recently discovered *D. platura* biotypes) has so far only been conducted by Savage et al. (2016) and Meraz-Álvarez et al. (2020). In some crops, members of the SMC can occasionally be more abundant than other specialized *Delia* pests although their relative abundance in relation to that of specialists shows a high degree of variation between studies and season (Doane & Chapman, 1962; Finlayson, 1956; Foley & Stone, 1958; Meraz-Álvarez et al., 2020; Merrill, 1951; Merrill & Hutson, 1953; Miles, 1948; Nair & McEwen, 1973; Salgado et al., 2026; Savage et al., 2016). For example, Nair & McEwen (1973) reported that while cabbage maggots (*D. radicum* (Linnaeus)) were dominant during most of the season in radish (*Raphanus sativus*) crops in Ontario, Canada, SMC members became more abundant late in the season. Conversely, in onion *(Allium cepa*) crops, SMC members have been found to be more abundant than the onion maggot (*D. antiqua* (Meigen)) early in the season in both the United Kingdom (Miles, 1948) and in Quebec, Canada (Fortier, 2021). Because the SMC is generally treated as a single unit rather than three distinct biological entities, there is generally little information available on the phenology and abundance of each species and biotype and whether they vary according to crop in the field.

In Québec, one of the few Canadian provinces where the distribution of the two *D. platura* biotypes overlap (Savage et al., 2016), crops are potentially under threat of infestation by all SMC members. The two *D. platura* biotypes have been shown to differ in their mating system (Bush-Beaupré et al., 2023) and are asymmetrically reproductively incompatible (Bush-Beaupré, et al., 2024). These biological differences, along with apparent differences in their relative abundance and phenology (Savage et al., 2016; Van Der Heyden et al., 2020) may bias the interpretation of most works published on *Delia platura* dynamics in Québec (e.g. Lamb & Boivin, 2018) and potentially elsewhere in Eastern North America (Cho et al., 2025; Olaya-Arenas et al., 2024). Consequently, the effectiveness of control methods developed in studies where all members of the SMC were pooled may be compromised and/or the timing of their applications may not be optimized.

Control measures for the SMC in vegetable crops rely mainly on the prophylactic application of insecticides at planting, either as in-furrow treatments or seed coatings (Salgado & Nault, 2023; Wilson et al., 2015) along with cultural practices such as avoiding fields with high decomposing organic matter or crop residues (Hammond, 1995) and sowing under environmental conditions that favor rapid germination (Throne & Eckenrode, 1985), which are not always feasible. To limit the negative impacts of synthetic insecticides, which are not as effective as previously thought (Labrie et al., 2020), it is imperative to develop targeted and sustainable control alternatives. The sterile insect technique (SIT) is one such alternative that has been successfully applied to other *Delia* species in Canada and elsewhere (Fortier et al., 2024, 2025; Ticheler et al., 1980) and efforts are currently deployed towards the development of a SIT program for the two *D. platura* biotypes (Fortier et al., 2026).

Based on three years of pooled larval data collected from various vegetable crops in southern Quebec, Canada, Van der Heyden et al. (2020) found that spring infestations of *D. platura* biotype N appeared in the field before those of biotype H. To determine the optimal timing for field releases of sterile males for these biotypes in selected crops, we are building here on the work of Van der Heyden et al. (2020) by describing the field abundance and phenological trends of each SMC members in broccoli (*Brassica oleracea*), green onion and dry onion for the Montérégie region of Quebec, Canada.

### Green and dry onion sampling

Each year, the agricultural consulting firm PRISME (Productions en Régie Intégrée du Sud de Montréal enr.) scouts over 130 dry onion and 60 green onion fields on 23 farms located in the Napierville county (MRC Des Jardins-de-Napierville, QC, Canada, 73°24’ to 73°37’ West, 45°10’ to 45°16’ North), totalling over 1 000 hectares. Damage incidence caused by *Delia* spp. is assessed once a week on one-meter sites (1.5 sites/acre) in each field. *Delia* larvae found in damaged plants were therefore collected opportunistically by field scouts during the 2017 to 2022 seasons, from early May to July (see Table 1). All larvae were examined morphologically to separate those of the onion maggot *D. antiqua* from those of the SMC following Savage et al. (2016). Members of the SMC were then identified to species and biotype using the High-Resolution Melting (HRM) PCR assay of Van der Heyden et al. (2020). We report here only the data pertaining to the SMC.

**Table 1.**
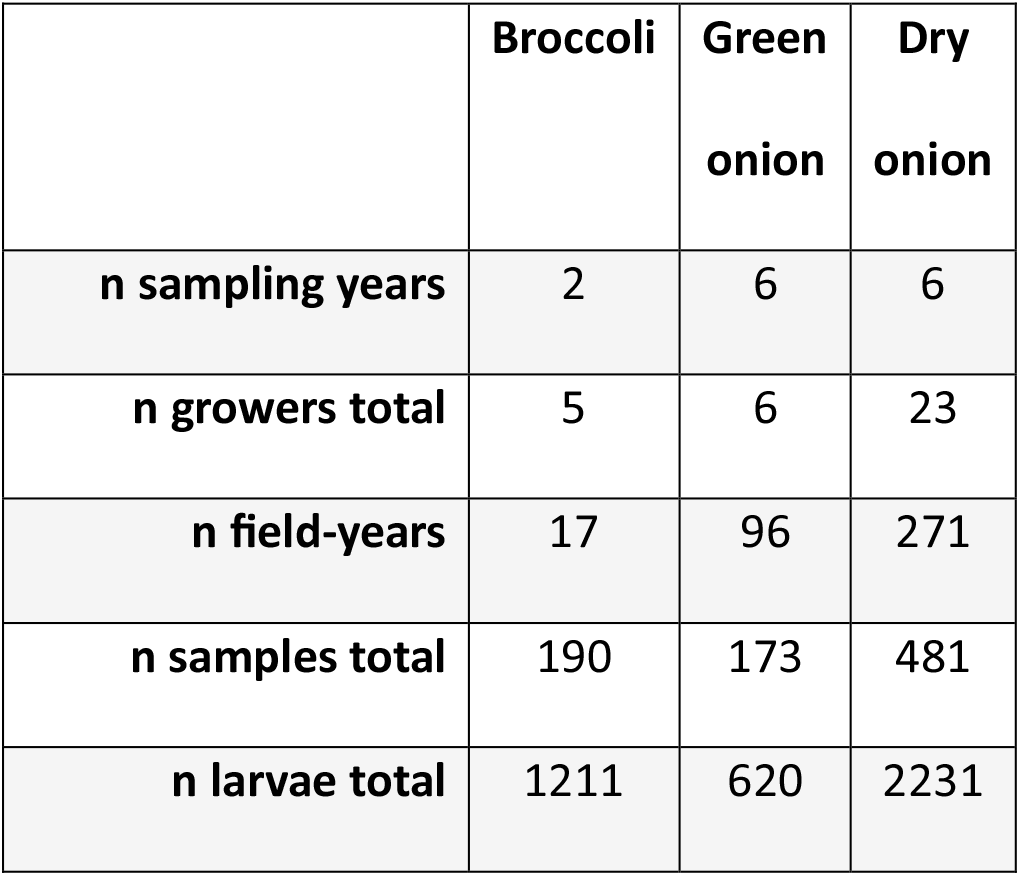
Number of years, growers, field-years, samples and larvae belonging to the seedcorn maggot complex *(Delia platura* biotypes H and N, *D. florilega* (Diptera: Anthomyiidae). Samples collected from broccoli (*Brassica oleracea*) and onion (*Allium cepa*) crops between 2017 and 2022 in western Montérégie, Québec, Canada. Field-years: sum of the number of year(s) each field was sampled. Samples: number of days for which at least one larva from unique fields were collected.

|  | Broccoli | Green<br>onion | Dry<br>onion |
| --- | --- | --- | --- |
| **n sampling years** | 2 | 6 | 6 |
| **n growers total** | 5 | 6 | 23 |
| **n field-years** | 17 | 96 | 271 |
| **n samples total** | 190 | 173 | 481 |
| **n larvae total** | 1211 | 620 | 2231 |

### Broccoli sampling

*Delia* larvae were systematically sampled in broccoli fields twice weekly in 2019 and 2020 from mid-May to September in western Montérégie, Québec. Sampling consisted of digging around the base of the plants to collect the larvae around the roots and base of the stem using tweezers. The sampling scheme consisted of 10 consecutive plants at 10 randomly selected sites per field. Some exceptions to the sampling scheme were made when many (> 50) larvae were collected on a given date such that the number of sites and consecutive plants were reduced. Identification procedure was the same as for onions except that larvae were first examined to separate those belonging to the cabbage maggot *D. radicum* from those of the SMC following Savage et al. (2016).

### Data analysis

Phenological and abundance data for each crop were analysed with separate Bayesian hierarchical generalized additive models with a negative binomial likelihood and log link function. A spline was fit to each SMC member as a function of day of year along with random intercepts for field, grower and year IDs. Models were fit using the brms library (v. 2.22.13; Bürkner, 2017) in R (v. 4.5.1; R Core Team, 2025). Model priors were chosen and tuned using prior predictive checks. Model diagnostics included posterior predictive checks and inspection of trace plots and R-hat values.

## Results

### Green and dry onion

Trends were similar for both dry and green onion (Figure 1). In these crops, *D. platura* biotype N was dominant during the month of May with a clear shift in early June when both *D. platura* biotype H and *D. florilega* became more abundant. These results are congruent with those of Van der Heyden et al. (2020), who found biotype N to appear approximately 17 days before biotype H. Thus, a sensible approach to managing the SMC in these two *Allium* crops may be to release sterile flies of biotype N before June and switch to biotype H afterwards. While *D. florilega* is not currently under evaluation for potential use in a SIT program, our results indicate that it must not be dismissed as a pest of onion crops since it was nearly as abundant as either *D. platura* biotype during certain time windows (Fig. 1).

**Figure 1.**
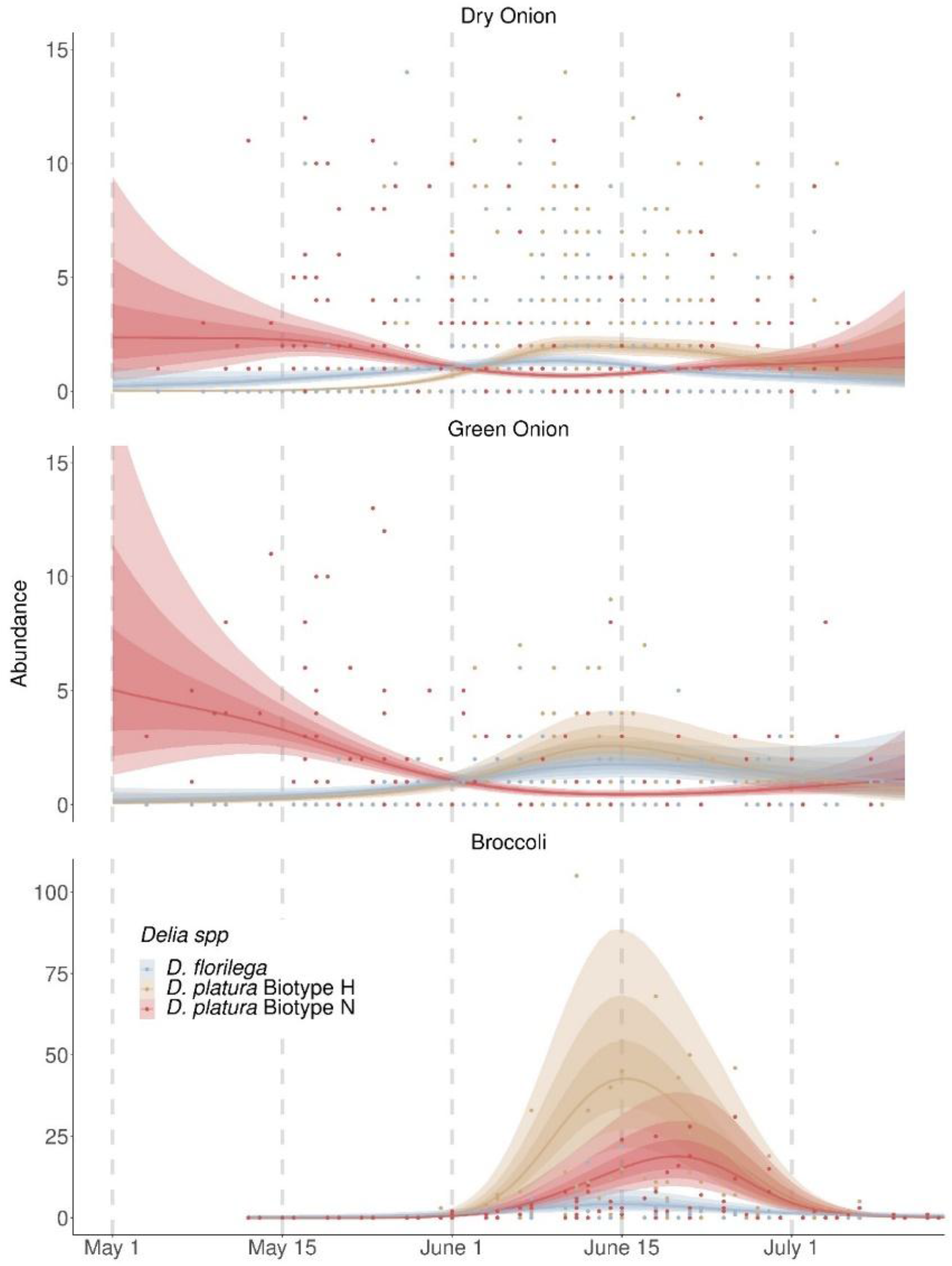
Mean abundance of SMC larvae in dry and green onion and broccoli. Line-intervals represent the median, 50%, 80% and 95% intervals of the posterior distribution of the mean abundance of each SMC member predicted by a Bayesian hierarchical generalized additive model.

### Broccoli

In broccoli fields, which were planted and not sown, no SMC larvae were found before the end of May (approximately two weeks after planting), and once they appeared, the three taxa showed considerable overlap during the month of June (Figure 1). All SMC members were observed within a similar same timeframe during the June abundance peak, but the abundance of *D. florilega* was considerably lower than that of the two *D. platura* biotypes. A second but more modest abundance peak was also observed during the first 2-3 weeks of August and while our sample size for that second peak was limited, we did not collect any *D. florilega* individuals over that period (Supplemental Figure 1). Our results indicate that broccoli is at higher risk of damage by both *D. platura* biotypes during the month of June, with biotype H appearing a few days earlier and in higher abundance. Sterile flies from both biotypes should therefore be released concurrently to reduce *D. platura* populations in broccoli.

## Conclusion

*Delia florilega* and *D. platura* biotype H displayed similar phenological trends in all sampled crops whereas biotype N showed highly contrasting trends between *Allium* and broccoli. The highest abundance of biotype N in *Allium* was in early May and tapered off towards the beginning of June but in broccoli, it was only found during the month of June (for the first peak). While multiple hypotheses may be posited to explain these contrasting phenologies, testing them was beyond the scope of the present work. Nonetheless, the phenological trends we report here are congruent with those of other works and emphasize the need to treat all three SMC members as separate biological entities when developing pest-specific control measures (Bush-Beaupré et al., 2023; Bush-Beaupré et al., 2024; Van Der Heyden et al., 2020). With the SIT currently being developed for the two *D. platura* biotypes (Fortier et al., 2026), it is imperative that the field release of sterile flies from each biotype be scheduled ahead of their larval activity window for each target crop.

## Acknowledgements

We are grateful to the participating farms for their trust and access to the fields and for all the field scouts and research assistants that participated in specimen collection. This project was made possible with funding from Agriculture and Agri-Food Canada (Canadian Agri-Science for Horticulture 3, #ASC18/19-Activity 8) and by the “Innov’Action agroalimentaire” program (grant number IA119539), offered under the Canadian Agricultural Partnership, an agreement between the Government of Canada and the Gouvernement du Québec with contributions from Compagnie de recherches Phytodata Inc., Productions en Régie Intégrée du Sud de Montréal Enr. (PRISME), and Bishop’s University.

## Supplemental material

**Supplemental Figure 1.**
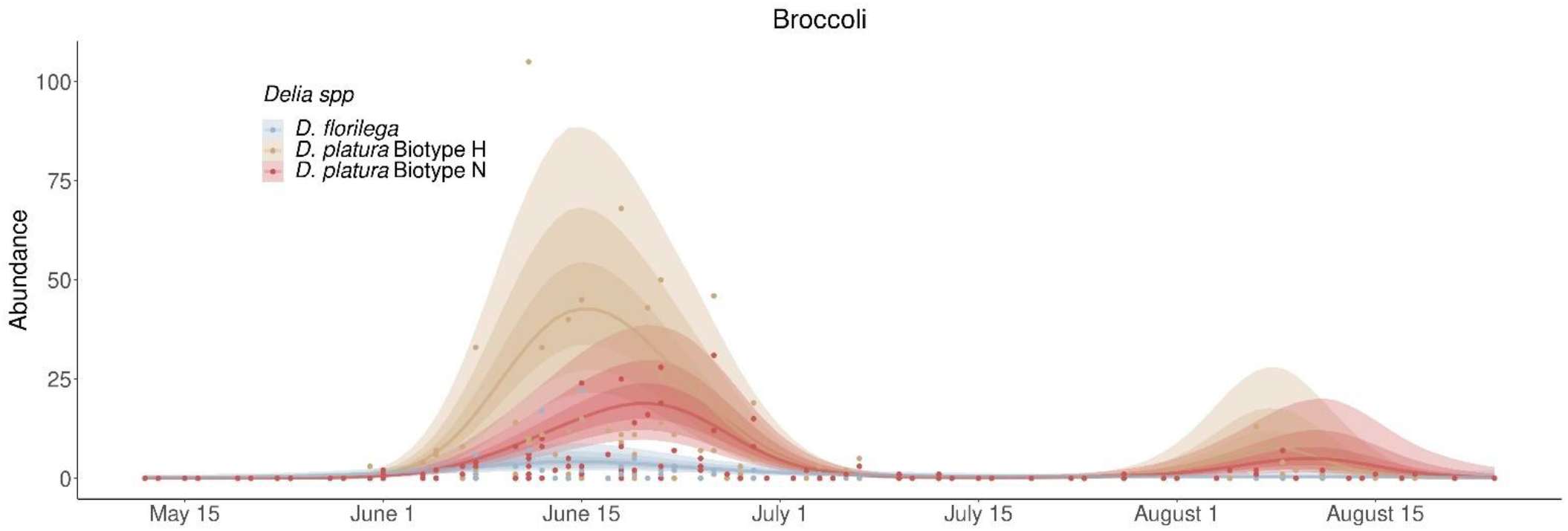
Mean abundance of the three SMC members for broccoli displaying the two abundance peaks observed. Line-intervals represent the median, 50%, 80% and 95% intervals of the posterior distribution of the mean abundance of each SMC member predicted by a Bayesian hierarchical generalized additive model.

**Supplemental Table 1.** Literature reports of relative importance of damage of *Delia* pest species. List of reports on the relative importance of damage caused by specialist and generalist *Delia* pest species in three crops from North America and United Kingdom. SMC = Seed Maggot Complex (*Delia platura* (Biotypes H and N) and *Delia florilega*).

| Crop | Geographical Location | *Delia spp* | Relative damage contribution | Reference |
| --- | --- | --- | --- | --- |
| Onion<br>(*Allium cepa*) | British Columbia, Canada | *D. antiqua*<br>SMC | SMC ≈ *D. antiqua* | (Finlayson, 1956) |
|  | England & Wales, U.K. |  | Whole season: *D. antiqua* > SMC<br>Early season: SMC > *D. antiqua* | (Miles, 1948) |
|  | Michigan, U.S.A. |  | *D. antiqua* > SMC | (Merrill, 1951; Merrill & Hutson, 1953) |
| | Quebec, Canada | | SMC ≈ *D. antiqua*<br>(Biotype N > Biotype H) | (Savage et al. 2016) |
|  | Great Lakes, North America |  | Whole season: *D. antiqua* > SMC<br>Early season: SMC > *D. antiqua* | (Salgado et al., 2026) |
|  | Eastern Oregon/western Idaho, USA |  | Whole season: *D. antiqua* > SMC<br>Early season: SMC > *D. antiqua* | (Salgado et al., 2026) |
| Broccoli | Quebec, Canada | *D. radicum*<br>SMC | SMC ≈ *D. radicum* | (Savage et al. 2016) |

| (Brassica<br>oleracea) |  |  | (Biotype H ><br>Biotype N) |  |
| --- | --- | --- | --- | --- |
|  | Mexico | *D. planipalpis*<br>*D. platura* | *D. planipalpis* ><br>*D. platura* | (Meraz-Álvarez et al.<br>2020) |

